# A ‘Naturalistic Neuroimaging Database’ for understanding the brain using ecological stimuli

**DOI:** 10.1101/2020.05.22.110817

**Authors:** Sarah Aliko, Jiawen Huang, Florin Gheorghiu, Stefanie Meliss, Jeremy I Skipper

**Affiliations:** London Interdisciplinary Biosciences Consortium, University College London, UK; Experimental Psychology, University College London, UK; School of Psychology and Clinical Language Sciences, University of Reading, UK

## Abstract

Neuroimaging has advanced our understanding of human psychology using reductionist stimuli that often do not resemble information the brain naturally encounters. It has improved our understanding of the network organization of the brain mostly through analyses of ‘resting-state’ data for which the functions of networks cannot be verifiably labelled. We make a ‘*Naturalistic Neuroimaging Database*’ (NNDb v1.0) publically available to allow for a more complete understanding of the brain under more ecological conditions during which networks can be labelled. Eighty-six participants underwent behavioural testing and watched one of 10 full-length movies while functional magnetic resonance imaging was acquired. Resulting timeseries data are shown to be of high quality, with good signal-to-noise ratio, few outliers and low movement. Data-driven functional analyses provide further evidence of data quality. They also demonstrate accurate timeseries/movie alignment and how movie annotations might be used to label networks. The NNDb can be used to answer questions previously unaddressed with standard neuroimaging approaches, progressing our knowledge of how the brain works in the real world.

## Background & Summary

A primary goal of human neuroscience is to understand how the brain supports broad psychological and cognitive functions that are engaged during everyday life. Progress towards achieving this goal over the last two decades has been made with tens of thousands of task- and resting-state based functional magnetic resonance imaging studies (henceforth, task-fMRI and resting-fMRI). While these studies have led to a number of important discoveries, there is mounting evidence that a better understanding of brain and behaviour might be achieved by also conducting studies with more ecologically valid stimuli and tasks (natural-fMRI).

### Task-fMRI

For task-fMRI, general psychological processes are decomposed into discrete (though hypothetical) component processes that can theoretically be associated with specific activity patterns. To ensure experimental control and because of reliance on the subtractive method^1^, these components are studied with stimuli that often do not resemble things participants might naturally encounter and tasks they might actually perform in the real-world (a topic long debated)^2–4^. For example, language comprehension has been broken down into component processes like phonology and semantics. These individual subprocesses are largely localised in the brain using isolated auditory-only ‘speech’ sounds (like ‘ba’) in the case of phonology and single written words in the case of semantics^5^. Participants usually make a meta-linguistic judgement about these stimuli, with a corresponding button response (e.g., a ‘2AFC’ indicating whether a sound is ‘ba’ or ‘pa’).

The result of relying on unnatural stimuli and tasks is that our neurobiological understanding derived from task-fMRI may not be representative of how the brain processes information. This is perhaps why fMRI test-retest reliability is low^6,7^. Indeed, more ecologically valid stimuli like movies have higher reliability than resting- or task-fMRI. This is not only because these enhance activity, decrease head movement and improve participant compliance^8,9^. Rather, natural stimuli have higher test-retest reliability mostly because they are more representative of operations the brain normally performs and provide more constraints on processing^10–15^.

### Resting-fMRI

There has arguably been a significant increase in our understanding of the network organization of the human brain because of the public availability of large resting-fMRI datasets, analysed with dynamic and other functional connectivity methods^16,17^. These include the INDI ‘1000 Functional Connectomes Project’^18^, ‘Human Connectome Project’ (HCP)^19^ and UK Biobank^20^. Collectively, these datasets have more than 6,500 participants sitting in a scanner ‘resting’. Resulting resting-state networks are said to represent the ‘intrinsic’ network architecture of the brain, i.e., networks that are present even in the absence of exogenous tasks. These networks are often claimed to be modular and to constrain the task-based architecture of the brain^21^.

As with task-fMRI, one might ask how representative resting-state networks are given that participants are anything but at rest. They are switching between fixating on a cross-hair, trying to stay awake, visualising, trying not to think and thinking through inner speech^21,22^. Some of these are not particularly natural and, unlike task-fMRI, there is no verifiable way to label resulting networks. At best, reverse inference is used to give 5-10 gross labels, like the ‘auditory’ network^23–25^. Despite claims that these ‘intrinsic’ networks constrain task-fMRI networks, it is increasingly suggested that this is not necessarily so^21^. The brain is less modular during task-compared to resting-fMRI^26^ and modularity decreases as tasks get more difficult^27–29^. Indeed, up to 76% of the connections between task- and resting-fMRI differ^30^. Furthermore, more ecological stimuli result in *new* sets of networks that are less modular and only partly constrained by resting networks^31,32^.

### Natural-fMRI

Based on considerations like these, there is a growing consensus that taking a more ecological approach to neuroscience might increase our understanding of the relationship between the brain and behaviour^5,33–42^. Though there are 16 publicly available natural-fMRI datasets that might be used for this purpose^43^, there is no dataset with a large number of participants, long and natural stimuli and stimulus variability. Specifically, most datasets have a small number of participants (median = 23). However, 80 or more participants are preferred for detecting medium effect sizes and producing replicable task-fMRI results^44–46^. Natural-fMRI datasets with larger numbers tend to use short (∼10 minute) audio or audiovisual clips. However, stimulation times of 90 minutes or more are preferred for reliability and individual analyses^47–50^.

Longer natural-fMRI datasets have a small number of participants and one stimulus (though see^51^). These include 11 people watching ‘Raiders of the Lost Ark’^52^ and 20 listening to an audio description of ‘Forrest Gump’ during fMRI. A subset of the latter returned to be scanned watching the movie dubbed in German (http://studyforrest.org)^53,54^. However, with only one movie, generalisability is limited. More movies would not only increase generalisability, they would increase the number of stimulus features and events in a variety of (jittered) contexts that might be annotated. These could then be used to label finer grained patterns of activity, e.g., making machine learning/decoding approaches more feasible^55–57^.

Indeed, there is no a priori reason participants need to watch the same movie during natural-fMRI. Existing long datasets might use one stimulus because intersubject correlation (ISC) is a common method for analysing natural-fMRI data^58^. Though this is a powerful model-free approach (for an overview, see^59^), it requires participants to watch the same movie. However, most questions are stimulus-feature or event specific and independent of the movie being watched. Thus, model-free independent component, convolution/deconvolution and other analyses can be done at the individual participant level with different movies, increasing generalisability and the possibility of more detailed analyses through more varied stimulus annotations.

### NNDb

To fill these gaps in publicly available data, we collected a ‘*Naturalistic Neuroimaging Database*’ (NNDb) from 86 people who each did a battery of behavioral tests and watched a full-length movie during natural-fMRI. We sought to reach a balance that promotes generalizability, allows a large variety of stimulus features and events to be annotated and most any analysis method to be used. To achieve this, our participants watched 10 different movies from 10 different genres. They had not previously seen the movies they watched because multiple viewings might change the functional network architecture of the brain (though activity patterns may appear similar)^60^. We validate that the data is of a high quality and good temporal alignment, whilst providing an example of using annotations to label networks. Figure 1 provides an overview of the study and analyses used to make this assessment.

**Figure 1.**
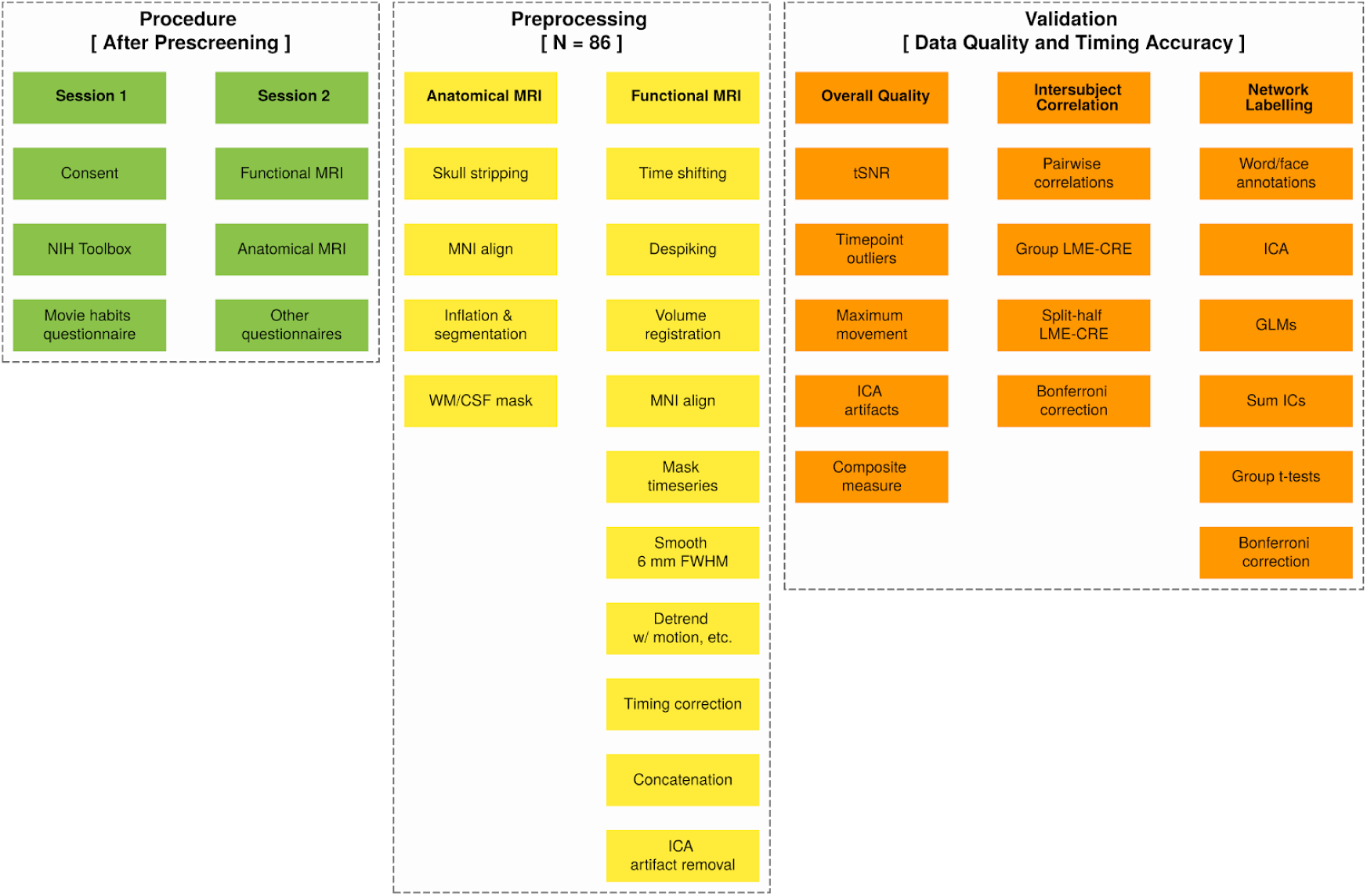
Schematic overview of NNDb study procedures, preprocessing and data validation. Procedures (green) occurred over two sessions separated by a few weeks. Session one consisted primarily of a battery of behavioural tests to quantify individual differences. In session two, functional magnetic resonance imaging (MRI) was done while participants watched one of 10 full length movies followed by anatomical MRI. After preprocessing the data (yellow), three primary data validation approaches were undertaken (orange). Functional MRI data is shown to be relatively free of outliers, with good temporal signal-to-noise ratio (tSNR) and low numbers of outlying timepoints, head movement and independent component analysis (ICA) artifacts (orange, column 1; see Tables 5-6 and Figure 2). Intersubject Correlation analyses provide evidence for functional data quality and the temporal synchronization between participants and movies using linear-mixed effects models with crossed random effects (MNE-CRE; orange, column 2; see Figure 3). Automated word and face annotations were used to find associated independent component (IC) timecourses from ICA using general linear models (GLMs; orange, column 3; see Figure 4). In addition to further illustrating data quality and timing accuracy, this analysis shows how annotations might be used to label brain networks.

Data discovery is nearly unlimited with the NNDb as there are a vast number of annotations that can be made from the movies and approaches to analysis. This flexibility makes it usable across disciplines to address questions pertaining to how the brain processes information naturally. This includes more than replicating prior findings with more ecologically valid stimuli. That is, there are a number of broad open questions that the NNDb can be used to address for the first time, like the systematic study of how context is used by the brain^5^. Given the lack of robust neuroimaging biomarkers for mental illness^61,62^, the NNDb might also help increase the pace of clinically relevant discovery, e.g., by uncovering labelled network patterns that predict individual differences^42^.

## Methods

### Participants

Based on sample size considerations reviewed above, we attempted to create a database with 84 participants watching 10 full-length movies from 10 genres (Table 1). To reach this number, we recruited 120 individuals using participant pool management software (http://www.sona-systems.com/). Those meeting MRI safety (e.g., no metal implants) and inclusion criteria were invited to participate. In particular, participants were required to be right-handed, native English speakers, with no history of claustrophobia, psychiatric or neurological illness, not taking medication (at the time of scan), without hearing impairment and with unimpaired or corrected vision. We also pseudo-randomly selected participants such that the final sample was relatively gender balanced. Of those who enrolled and completed the study, two were excluded as they were determined to be left handed, two because they asked to get out of the scanner multiple times and one who was a data quality outlier.

**Table 1.**
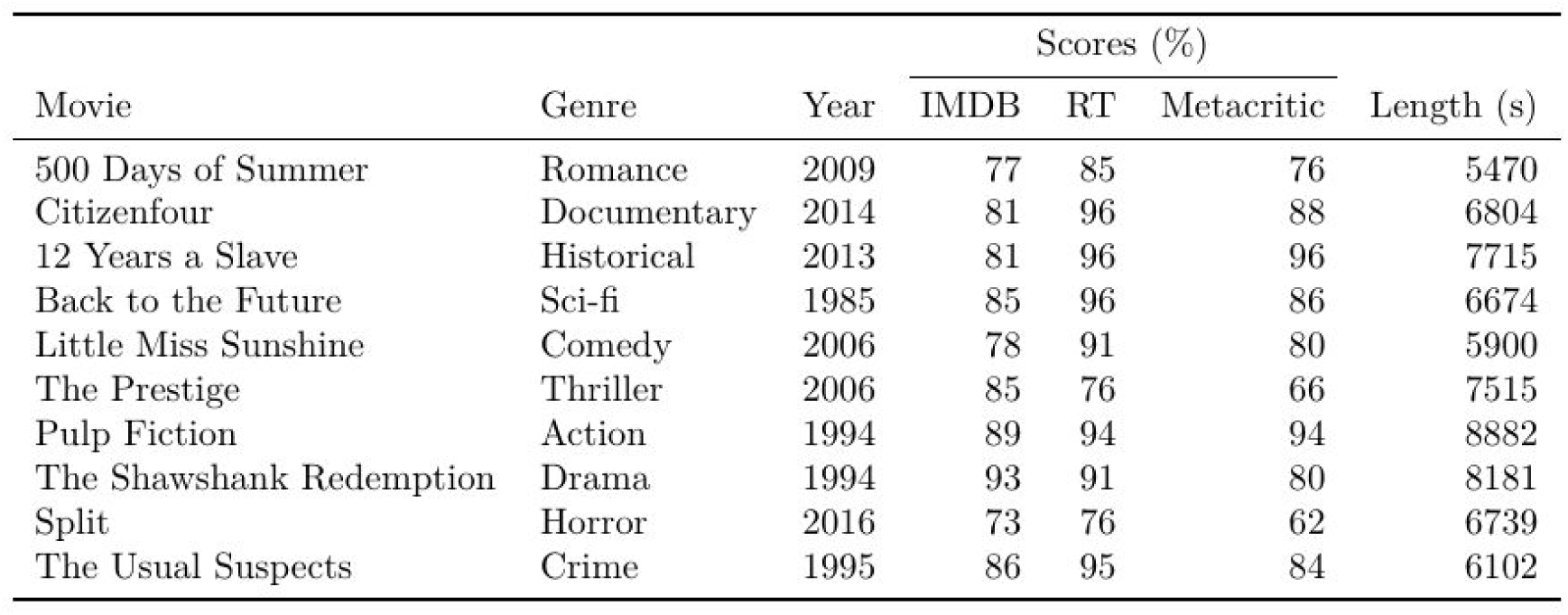
Description of the movies used in the *naturalistic neuroimaging database*. Ten full length movies were chosen from 10 genres. These were required to have been successful, defined as an average *Internet Movie Database* (IMDb), *Rotten Tomatoes* (RT) and *Metacritic* score greater than 70%. IMDb scores were converted to percentages for this calculation. Movie lengths are given in seconds (s), also equivalent to the number of whole brain volumes collected when participants watched these movies during functional magnetic resonance imaging.

The final sample consisted of 86 participants (42 females, range of age 18–58 years, M = 26.81, SD = 10.09 years). These were pseudo-randomly assigned to a movie they had not previously seen, (usually) from a genre they reported to be less familiar with. Table 2 provides a summary of participant demographics by movie. At the conclusion of the study, participants were given £7.5 per hour for behavioural testing and £10 per hour for scanning to compensate for their time (receiving ∼£40 in total). The study was approved by the ethics committee of University College London and participants provided written informed consent to take part in the study and share their anonymised data.

**Table 2.** Description of participants in the *naturalistic neuroimaging database*. All participants (N) were right-handed and native English speakers. Gender is expressed as percent female. Ethnic diversity is expressed as percent Black, Asian and Minority Ethnic (BAME). Educational attainment is expressed as percent with a Bachelor’s degree or higher. Data roughly match 2011 London, UK consensus data (https://data.london.gov.uk/census/). We include the ‘Cognition Fluid Composite v1.1’ and ‘Negative Affect Summary (18+)’ ‘T-scores’ as example tests from the National Institute of Health (NIH) Toolbox battery. The bottom two rows are the means and standard deviations of row means weighted by number of participants (wMeans/wSD).

| N | Movie | Age | Female (%) | BAME (%) | Monolingual (%) | ≥ Bachelor's (%) | | NIH Toolbox Examples | |
| --- | --- | --- | --- | --- | --- | --- | --- | --- | --- |
|  |  |  |  |  |  | Participant | Mother | Fluid Cog. | Neg. Affect |
| 20 | 500 Days of Summer | 27.70 | 50.00 | 30.00 | 85.00 | 75.00 | 60.00 | 58.11 | 51.25 |
| 18 | Citizenfour | 27.00 | 50.00 | 41.18 | 77.78 | 61.11 | 66.67 | 53.71 | 51.22 |
| 6 | 12 Years a Slave | 27.17 | 50.00 | 66.67 | 50.00 | 50.00 | 50.00 | 31.00 | 50.83 |
| 6 | Back to the Future | 22.17 | 50.00 | 40.00 | 66.67 | 66.67 | 83.33 | 39.67 | 57.67 |
| 6 | Little Miss Sunshine | 35.67 | 33.33 | 66.67 | 66.67 | 50.00 | 16.67 | 45.67 | 52.33 |
| 6 | The Prestige | 37.00 | 50.00 | 0.00 | 100.00 | 83.33 | 33.33 | 76.00 | 51.67 |
| 6 | Pulp Fiction | 22.67 | 50.00 | 83.33 | 66.67 | 33.33 | 0.00 | 51.00 | 57.67 |
| 6 | The Shawshank Redemption | 22.17 | 50.00 | 100.00 | 50.00 | 50.00 | 83.33 | 68.00 | 50.00 |
| 6 | Split | 22.67 | 50.00 | 50.00 | 83.33 | 66.67 | 33.33 | 52.67 | 57.50 |
| 6 | The Usual Suspects | 23.17 | 50.00 | 66.67 | 83.33 | 100.00 | 33.33 | 55.33 | 54.50 |
| **86** | **wMean** | **26.93** | **48.84** | **48.62** | **75.58** | **65.12** | **51.16** | **54.01** | **52.79** |
|  | **wSD** | **4.41** | **4.27** | **24.62** | **13.48** | **15.99** | **23.49** | **10.47** | **2.67** |

### Procedures

Participants meeting inclusion criteria were scheduled for two sessions on seperate days. During session one, participants gave informed consent and then completed the majority of the National Institute of Health (NIH) Toolbox. This provides validated measures of sensory, motor, cognitive and emotional processing that might be used as individual difference measures^63^. We only excluded tests in the ‘Sensation’ and ‘Motor’ domains that required physical implementation (e.g., scratch and sniff cards, a pegboard, endurance walking, etc.). Participants were provided with headphones and tests were administered in a sound shielded testing room on an iPad. At the end of session one, participants filled out a questionnaire on movie habits, including information on preferred movie genres. The entire session typically took about one hour.

Functional and anatomical MRI and a final questionnaire were done during a second session that was separated from the first by about 2-4 weeks (M = 20.36 days; SD = 23.20). Once in the scanning suite, participants reporting corrected vision were fitted with MRI-safe glasses. All participants were fit with earbuds for the noise-attenuating headphones. They were put in the head-coil with pillows around the head and under the knees for comfort and to reduce movement over the scanning session. Once in place, participants chose an optimal stimulus volume by determining a level that was loud but comfortable. Video presentation was adjusted for optimal viewing quality. Participants were given a bulb in their right hand and told to squeeze if something was wrong or they needed a break during the movie. They were instructed to not move as best as they could throughout scanning as movement would make the scans unusable.

Except in one case, functional MRI movie scans were done first and with as few breaks as possible. During breaks, participants were told that they could relax but not move. During scanning, participants were monitored by a camera over their left eye. If they appeared drowsy or seemed to move too much during the movie, the operator of the scanner gave them a warning over the intercom by producing a beep or speaking to them. In some cases we stopped the scan to discuss with the participant. After the movie, participants had an anatomical scan and were told they could close their eyes if they wished. Following scanning, participants filled out other questionnaires, e.g., about their specific experience with content in the movie they watched. Finally, participants were paid and sent home.

### Movie Stimuli

Table 1 provides an overview of the 10 movies participants watched during fMRI. These were chosen to be from 10 different genres and to have an average score of > 70% on publicly available metrics of success. These were the *Internet Movie Database* (IMDb; https://www.imdb.com/), *Rotten Tomatoes* (RT; https://www.rottentomatoes.com/) and *Metacritic* (https://www.metacritic.com/). All movies were purchased and stored as ‘.iso’ files. Relevant sections of the DVD (i.e., excluding menus and extra feature) were directly concatenated to a mpg container using:

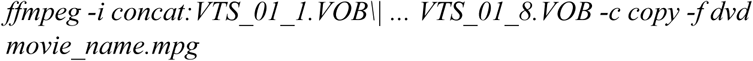

Where ‘-c’ copies the codec and ‘-f’ specifies the DVD format. This maintains the original DVD video size and quality.

The resulting files were presented to participants in full-screen mode through a mirror reversing LCD projector to a rear-projection screen measuring 22.5 cm x 42 cm with a field of view angle of 19.0°. This screen was positioned behind the head coil within the MRI bore and was viewed through a mirror above participants’ eyes. High quality audio was presented in stereo via a Yamaha amplifier through Sensimetrics S14 scanner noise-attenuating insert earphones (https://www.sens.com/products/model-s14/).

#### Movie Pausing

Movies were played with as few breaks as possible. This allows for the most natural, uninterrupted viewing experience. It also results in good timing accuracy, needed for relating movie features and events to brain responses. It maintains timing by avoiding unknown and accumulated human and hardware processing delays associated with starting and stopping and minimises discontinuities in the hemodynamic response. To accomplish continuous play with the possibility of arbitrary stopping points, we created a script and hardware device to allow the operator to stop the scanner and pause the movie at any time and resume where the movie left off when the scanner was restarted. Unless participants signalled that they wanted a break, the movies were played in about one hour segments (because of a software limitation on the EPI sequence we used). These breaks were timed to occur during scenes without dialogue or relevant plot action.

Specifically, a Linux BASH script opened and paused movies using ‘*MPlayer*’ (http://www.mplayerhq.hu/). The script then went into a state of waiting for a TTL (transistor-transistor logic) pulse from the scanner, indicating that scanning had begun. Pulses were received through a USB port connected to an Arduino Nano built to read and pass TTL pulses from the scanner to the script. When the scan was started and the first TTL pulse was received, eight seconds were allowed to elapse before the movie began to play. These timepoints allowed for the scanner to reach a state of equilibrium and were later discarded. If the scanner was subsequently paused, e.g., because the participant requested a break, the movie pausing BASH script stopped the movie within 100 ms. This known delay occurred because the script monitors for TTL pulses every 50 ms. If a pulse was not registered, the script required that the next pulse also did not arrive before pausing to assure pulses had stopped. When the scan was restarted, eight seconds were again allowed to pass before the movie was unpaused.

Whenever a movie was paused after it had been playing, the whole brain volume being collected was dropped, causing up to one second of the movie to be lost from the fMRI timeseries. There were two versions of the script. In the first, the movie picked up where it left off when it had been paused (v1; N = 29 or 33.72% of participants). The second version rewound the movie to account for the time lost from the dropped volume. However, because of an initial coding error in version two (v2.1), the script fast forwarded instead of rewound in N = 13 or 15.12% of participants. Because fast forwarding could not be greater than one second and the error affected only 47.44% of the runs for those 13 participants (with the other 52.56% being correctly rewound), data timing quality was not compromised more than the first version of the script on average. The error was subsequently fixed for the remainder of the study (v2.2; N = 44 or 51.16% of participants). In all versions, output files from the script recorded system and movie timing to calculate start, stop and rewind times. For this reason, all (including system) delays were tracked and are, therefore, known quantities that can be accounted for in preprocessing to assure that fMRI timeseries and movies are temporally well aligned (see ‘Timing correction’ below). The Supplementary Materials provides additional information on how the script kept track of timing information and rewind times were calculated.

#### Movie Annotations

Words and faces were annotated in the movies using fully automated approaches. These were then used to demonstrate data and timing quality while also illustrating a method for network labelling. For words, we extracted the audio track as a ‘.wav’ and the subtitle track as a ‘.txt’ file from each movie ‘.iso’ file. The .wav file was input into the ‘*Amazon Transcribe*’, a machine learning based speech-to-text transcription tool from *Amazon Web Services* (AWS; https://aws.amazon.com/transcribe/). The resulting transcripts contained on and offset timings for individual words, although not all words are transcribed or accurately transcribed. In contrast, movie subtitles do not have accurate on and offset times for individual words though most words are accurately transcribed. Therefore, to estimate the on and offset times of the words not transcribed, a script was written that first uses dynamic time warping (DTW^64^) to align word onsets from the speech-to-text transcript to corresponding subtitle words in each individual subtitle page. Subtitle words that matched or that were similar to the transcriptions during the DTW procedure inherited the timing of the transcriptions.

Remaining subtitle words not temporally labeled were then estimated, with different degrees of accuracy. Continuous and partial word estimations inherited their on and offset times from matching/similar transcription words in the subtitle page. ‘Continuous’ words use the on and offset times from adjacent words directly, making them the most accurate. ‘Partial’ estimation occured where there was more than one word between matched/similar words. In those cases the length of each word was approximated, making it less accurate. ‘Full’ estimation was the least accurate, occurring when there were no matching/similar words transcribed, and the onsets and lengths of the words were estimated from the onset and offset of the subtitle page. For partial and full estimations, word length was determined by counting the number of letters in each word and dividing up the bounding time proportionally. For example, if there were two words with 10 and five letters, they got 66.67% and 33.33% of the time, respectively. This procedure might occasionally result in unreasonably long word length estimations (e.g., because of a long dramatic pause between words). In such cases, we used a word truncation algorithm based on mean word lengths in conversational speech^65^. A more detailed account of how this script works is available in the Supplementary Materials.

We used the AWS ‘*Amazon Rekognition*’ application programming interface (API) to obtain machine learning based faces annotations (https://aws.amazon.com/rekognition/). To do this, the original ‘.mpg’ video files were first converted to ‘.mp4’ to have a H264 codec compatible with Amazon’s Rekognition guidelines. A script called the face recognition API without any special configuration or modification and the output was a ‘.json’ file. This contained timestamps every 200 ms, if a face was present, other details about the face (e.g. predicted age range, gender, position on screen and whether the mouth was open) and confidence levels.

### Acquisition

Functional and anatomical images were acquired on a 1.5T Siemens MAGNETOM Avanto with a 32 channel head coil (Siemens Healthcare, Erlangen, Germany). We used multiband EPI^66,67^ (TR = 1 s, TE = 54.8 ms, flip angle of 75°, 40 interleaved slices, resolution = 3.2 mm isotropic), with 4x multiband factor and no in-plane acceleration; to reduce cross-slice aliasing^68^, the ‘leak block’ option was enabled^69^. Slices were manually obliqued to include the entire brain. A slice or at most a few slices of the most inferior aspect of the cerebellum was occasionally missed in individuals with large heads. This EPI sequence had a software limitation of one hour of consecutive scanning, meaning each movie had at least one break. From 5,470 to 8,882 volumes were collected per participant depending on which movie was watched (see Table 1). A 10 min high-resolution T1-weighted MPRAGE anatomical MRI scan followed the functional scans (TR = 2.73 s, TE = 3.57 ms, 176 sagittal slices, resolution = 1.0 mm)^3^.

### Preprocessing

MRI data files were converted from IMA to NIfTI format and preprocessed to demonstrate data quality using mostly the AFNI software suite^70^. Individual AFNI programs are indicated parenthetically in subsequent descriptions.

#### Anatomical

The anatomical/structural MRI scan was corrected for image intensity non-uniformity (‘*3dUniformize*’) and deskulled using *ROBEX*^*71*^ in all cases except for one participant where ‘*3dSkullStrip*’ performed better. The resulting anatomical image was nonlinearly aligned (using ‘*auto_warp.py*’) to the MNI N27 template brain, an average of 27 anatomical scans from a single participant (‘Colin’)^72^. The anatomical scan was inflated and registered with Freesurfer software using ‘*recon-all*’ and default parameters (version 6.0, http://www.freesurfer.net)^73,74^. This served to create white matter and ventricle (i.e., cerebral spinal fluid containing) regions of interest that could be used as noise regressors. These regions were resampled into functional dimensions and eroded to assure they did not impinge on grey matter voxels. Finally, anatomical images were ‘defaced’ for anonymity (https://github.com/poldracklab/pydeface).

#### Functional

The fMRI timeseries were corrected for slice-timing differences (‘*3dTshift*’) and despiked (‘*3dDespike*’). Next, volume registration was done by aligning each timepoint to the mean functional image of the centre timeseries (‘*3dvolreg*’). For 23 (or 26.74%) of participants, localiser scans were redone because, e.g., the participant moved during a break and the top slice of the brain was lost. For these participants, we resampled all functional grids to have the same x/y/z extent (‘*3dresample*’) and manually nudged runs to be closer together (to aid in volume registration). For all participants, we then aligned the functional data to the anatomical images (‘*align_epi_anat.py*’). Occasionally, the volume registration and/or this step failed as determined by manual inspection of all data. In those instances we either performed the same procedure as for the re-localised participants (N = 5 or 5.81%) or reran the ‘*align_epi_anat.py*’ script, allowing for greater maximal movement (N = 6 or 6.98%). Finally, the volume-registered and anatomically-aligned functional data were (nonlinearly) aligned to the MNI template brain (‘*3dNwarpApply*’).

Next, we cleaned the timeseries, resulting in what we henceforth refer to as the ‘detrended timeseries’ for each run. Specifically, we first spatially smoothed all timeseries to achieve a level of 6mm full-width half maximum, regardless of the smoothness it had on input (‘*3dBlurToFWHM*’^75^). These were then normalised to have a sum of squares of one and detrended (‘*3dTproject*’) with a set of commonly used regressors^76^: These were 1) Legendre polynomials whose degree varied with run lengths (following a formula of [number of timepoints * TR]/150); 2) Six demeaned motion regressors from the volume registration; 3) A demeaned white matter activity regressor from the averaged timeseries in white matter regions; and 4) A demeaned cerebrospinal fluid regressor from the averaged timeseries activity in ventricular regions.

#### Timing Correction

To use stimulus annotations, timing correction was done to account for delays caused by the movie pausing script to assure that fMRI timeseries and movies are well aligned. Specifically, this script introduced a known 100 ms delay that was cumulative for each break in the movie. Furthermore, depending on the versions of the script, there was also a possible additional (cumulative) delay from not rewinding (v1) or mistakenly fastforwarding (v2.1). These delays were calculated from script output files created for this purpose. Furthermore, these files allowed us to quantify potentially variable soft and hardware delays and account for these as well. In particular, every voxel of the detrended timeseries was shifted back in time using interpolation to account for all delays, in the same manner as in slice timing correction but over all voxels uniformly (‘*3dTshift*’). Detailed information on how delays were calculated and applied are provided in the Supplementary Materials.

#### ICA Artifact Removal

Spatial independent component analysis (ICA) is a powerful tool for detecting and removing artifacts that substantially improves signal-to-noise ratio in natural-fMRI data^77^. First, we concatenated all detrended timeseries after timing correction. As in the HCP, we did spatial ICA on this timeseries with 250 dimensions using ‘*melodic*’ (version 3.14) from FSL^78^. Next, we labelled and removed artifacts from timeseries, following an existing guide for manual classification^79^. One of three trained authors went through all 250 components and associated timecourses, labelling the components as ‘good’, ‘maybe’, or ‘artifact’. As described in Griffanti et al.^79^, there are a typical set of ‘artifact’ components with identifiable topologies that can be categorised as ‘motion’, ‘veins’, ‘arteries’, ‘cerebrospinal fluid pulsation’, ‘fluctuations in subependymal and transmedullary veins’ (i.e., ‘white matter’), ‘susceptibility artefacts’, ‘multi-band acceleration’ and ‘MRI-related’ artefacts. Our strategy was to preserve signal by not removing components classified as ‘maybe’. On a subset of 50 datasets (58.14% of the data), a second author classified all components to check for consistency. The authors discussed discrepancies and modified labels as warranted. It was expected that, similar to prior studies, about 70-90% of the 250 components would be classified as artifacts^79^. Once done, we regressed the ICA artifact component timecources out of the detrended and concatenated timeseries (‘*3dTproject*’).

### Analyses

We used the preprocessed, detrended and concatenated timeseries with ICA-based artifacts removed (henceforth ‘fully detrended timeseries’) for several analyses meant to validate data quality. These included calculating the temporal signal-to-noise (tSNR) ratio as one of a set of metrics and a composite measure to assess data quality at the timeseries level (Overall Data Quality). We also did two whole-brain functional analyses using two previously established data-driven methods. One was intersubject correlation (ISC) analysis and the other involved labelling functional networks with annotations (Network labelling). These serve to show data quality similar to past work and provide evidence for timing accuracy between fMRI timeseries for participants and movies. The latter is crucial as movie breaks varied across participants, resulting in a small amount of temporal interpolation and psychological discontinuity across runs.

#### Temporal Signal-to-Noise Ratio

We calculated tSNR both before and after extensive preprocessing to demonstrate data quality and how it might improve after timeseries cleaning and artifact removal. Temporal SNR can be defined as the mean signal divided by the standard deviation of the signal over voxel timeseries^80^. Though multiband acceleration greater than one improves sensitivity over multiband one^68^, average multiband four tSNR tends to be between 40-60, lower than unaccelerated sequences^68,78^. A natural-fMRI dataset showed that manual ICA-based artifact rejection increased tSNR around 50 units, though this was not multiband data^77^. HCP multiband four tSNR increased by 30 after ICA cleanup of resting-state data^78^. Thus, we would expect to see a similar baseline level and improvement after ICA artifact removal. It is worth noting that unlike most other datasets, we have over 1.5 hours of data per participant, likely sufficient at those tSNR values for detecting effects sizes of 1% or less^81^.

We first calculated tSNR (‘*3dTstat*’) on three timeseries: 1) A minimally preprocessed timeseries that was corrected for slice timing, despiked, volume-registered and aligned to the anatomical image, timing-corrected and concatenated; 2) The same timeseries but blurred with a 6 mm FWHM (‘*3dBlurToFWHM*’); and 3) A fully preprocessed timeseries, detrended using white matter, ventricular, motion and ICA artifact timecourse regressors (‘*3dTproject*’). We then calculated mean tSNR for all three timeseries using a mask that included grey matter, with most white matter and ventricle voxels removed. We calculated effect sizes at a voxel level using:

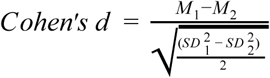

#### Overall Data Quality

Timeseries data quality were globally assessed using 10 measures and a composite of these: 1) To quantify timeseries timepoints outliers, we labelled voxels that were far from the median absolute deviation (‘*3dToutcount*’). Whole timepoints were defined as outliers if more than 10% of voxels were outliers by this definition; 2-8) We used seven parameters to quantify motion. These included the maximum average motion for each run from the demeaned motion regressors and the largest change in displacement between two successive timepoints (Delta); 9) The mean tSNR from the minimally preprocessed timeseries; and 10) The total number of ‘artifact’ ICA components. We then used the ‘*multicon*’ package in R (https://www.r-project.org/) to z-transform these 10 items and create a composite data quality score for each participant. We defined outlying participants as anyone whose composite score was more than three standard deviations from the mean.

#### Intersubject Correlation

In addition to illustrating data quality similar to prior results, ISC demonstrates synchrony of fMRI timeseries between our participants and movies. We compared the ISCs of participants watching the same movie to those watching different movies because a fundamental assumption of ISC is that synchrony is stimulus driven. Thus, we expected correlation values to be significantly greater for the same movie compared to different movies, with values similar to past ISC results from a large number of participants. For example, in a task-fMRI study with 130 participants, the maximum ISC is 0.27^82^.

Because movies had different lengths, we first truncated the fully detrended timeseries to be the length of the movie with the shortest duration (i.e., ‘*500 Days of Summer*’; 5470 s/TRs or about 1 hour and 31 minutes). We then computed pairwise Pearson’s correlations between the timeseries in each voxel for all pairs of participants for all movies (‘*3dTcorrelate*’). This resulted in (½ * 86 * (86-1)) = 3655 pairwise correlation maps. These are composed of (½ * 20 * (20-1))+(½ * 18 * (18-1))+((½ * 6 * (6-1)) * 8) = 463 maps from participants watching the same movie. The remaining (3655 - 463) = 3192 maps are from participants watching different movies.

For the group analysis, we first converted Pearson’s r values to be normally distributed as z-scores using the Fisher z-transformation. Then, to compare ISC maps from people watching the same or different movies, we used voxel-wise linear mixed effects models with crossed random effects (‘*3dISC*’). This approach accounts for the interrelatedness of the pairwise ISC maps and can handle unequal sample sizes^83^. The resulting map was Bonferroni-corrected for multiple comparisons using *t* = 6.04 corresponding to a voxel-wise p-value of .01 divided by the number of tests done in each voxel, i.e., *p* < .01/(4 * 64,542) = .00000004. We combined this with an arbitrary cluster size threshold of 20 voxels. To demonstrate reliability, we also repeated this analysis after splitting the data into groups of participants watching two different sets of five movies. We compared the resulting spatial patterns of activity using correlation in the unthresholded data and the Dice coefficient for thresholded data (*‘3ddot’*).

#### Network Labelling

Besides demonstrating data and timing quality, here we also illustrate a fairly straightforward method for using annotations to label networks with a method similar to one used in existing natural-fMRI studies. This combines model-free ICA to find networks and a model-based approach to label those networks^84,85^. In particular, we derive networks in each participant with ICA using ‘*melodic*’ run on the fully detrended timeseries (and, again, limited to 250 dimensions). We then convolve annotated word, no word, face and no face onsets and durations with a canonical hemodynamic response function (‘*3dDeconvolve*’). The resulting ideal waveforms are regressed against the 250 independent component timescourses using general linear model (GLMs) followed by pairwise contrasts between words and no words and faces and no faces (using FSL’s ‘*fsl_glm*’). A Bonferroni-corrected threshold was set at p = .01 at the single voxel level divided by 250 components and eight statistical tests (not all of which are discussed here), i.e., .01/(250 * 8) = p < .000005. We combined this with an arbitrary cluster size threshold of 20 voxels at the component level. If there was more than one resulting component at this threshold and cluster size, we summed those components.

For group analysis, we did one sample t-tests for GLM results of words vs no words, no words vs words, face vs no faces and no faces vs faces (‘*3dttest*++’). To correct for multiple comparisons, we again used a Bonforoni correction of .01 at the single voxel level divided by approximately 85,000 voxels and four tests, i.e., .01/(85,000 * 4), rounding to p < .00000001. We again combined this with an arbitrary cluster size threshold of 20 voxels. To illustrate the precise anatomical correspondence of our results with prior data, we overlay fMRI term-based meta-analysis from Neurosynth^86^ (Retrieved May 2020) for ‘language’ (https://neurosynth.org/analyses/terms/language/; from 1101 studies) and the ‘fusiform face’ area (https://neurosynth.org/analyses/terms/fusiform%20face/; from 143 studies; FFA). We further illustrate anatomical correspondence by showing the mean peaks of the putative (left and right) FFA, derived by averaging peaks from a meta-analysis of 49 studies (converted to MNI x/y/x coordinates = 39/-53/-22 and -40/-54/-23; see Table 1 in^87^).

## Data Records

Information and anatomical data that could be used to identify participants has been removed from all records. Resulting files are available from the OpenNeuro platform for sharing fMRI (and other neuroimaging) data at https://openneuro.org/datasets/ds002837 (dataset accession number: ds002837). A README file there provides a detailed description of the available content. Additional material and information are available on the NNDb website at http://www.naturalistic-neuroimaging-database.org.

### Participant Responses

**Location** demographics.csv

**File format** comma-separated value

Participants’ responses to demographic questions, the NIH Toolbox and all other questionnaires are available in a comma-separated value (CSV) file. Data is structured as one line per participant with all questions and test items as columns.

### Anatomical MRI

**Location** sub-<ID>/anat/sub-<ID>_T1w.nii.gz

**File format** NIfTI, gzip-compressed

**Sequence protocol** sub-<ID>/anat/sub-<ID>_T1w.json

The defaced raw high-resolution anatomical images are available as a 3D image file, stored as sub-<ID>_T1w.nii.gz.

The N27 MNI template aligned anatomical image and the anatomical mask with white matter and ventricles eroded are also available as

derivatives/sub<ID>/anat/sub-<ID>_T1w_MNIalignment.nii.gz and

derivatives/sub<ID>/anat/sub-<ID>_T1w_mask.nii.gz respectively

### Functional MRI

**Location** sub-<ID>/func/sub-<ID>_task-[movie]_run-0[1–6]_bold.nii.gz

**Task-Name [movie]** 500daysofsummer, citizenfour, theusualsuspects, pulpfiction, theshawshankredemption, theprestige, backtothefuture, split, littlemisssunshine, 12yearsaslave

**File format** NIfTI, gzip-compressed

**Sequence protocol** sub-<ID>/func/sub-<ID>_task-[movie]_run-0[1–6]_bold.json

Functional MRI data are available as individual timeseries files, stored as sub-<ID>_task-[movie]_run-0[1–6]_bold.nii.gz. The fully detrended timeseries is also available as derivatives/sub-<ID>_task-[movie]_bold_preprocessedICA.nii.gz.

### Motion and Outlier Estimates

**Location** motion/sub<ID>/sub-<ID>_task-[movie]_run0[1-6]_bold_[estimates].1D

**Motion [estimates]** motion, maxdisp_delt, wm, ventricle, outliers

**File format** plain text

Motion estimates are from the registration procedure in the AFNI program ‘*3dvolreg*’ and outliers were estimated using ‘*3dToutcount*’, These are provided in space-delimited text files where the estimates represent 1) *motion*: degree of roll, pitch and yaw and displacement in the superior (dS), left (dL) and posterior (dP) directions in mm; 2) *maxdisp_delt*: maximum displacement (delta) between any two consecutive timepoints; 3) *wm*: mean activity in the white matter; 4) *ventricle*: mean activity in the ventricles and 5) *outliers*: individual timepoint outliers at 10% levels.

### ICA Artifact Labels

**Location** derivatives/sub<ID>/func/sub-<ID>_task-[movie]_bold_ICAartifacts.1D

**File format** plain text

ICA components labeled as artifacts used to correct ICA time series as proved are provided as space delimited text where the columns are artifactual timecources.

### Annotations

**Location** stimulus/task-[movie]_[annot]-annotation.1D

Annotation [annot] word, face

**File format** plain text

Word, no word and face and no face onsets and durations are provided in four space-delimited text files. In the word annotation file, columns represent: 1) Words; 2) Word onset in seconds and milliseconds; 3) Word offset in seconds and milliseconds. In the face annotation file, columns represent: 1) Face onset in seconds and milliseconds and 2) Duration of face presence in seconds and milliseconds.

## Technical Validation

### Stimuli

#### Timing

The movies were played in the original DVD audio and video quality. This relative lack of compression results in low latencies when starting and stopping the movies. System delays were calculated from the timing output of the movie-pausing script. Averaged over all runs and participants, this delay was 19.73 ms (SD = 7.57). This is perhaps not more than the expected latency on a standard Linux kernel^88^. However, because this delay was measured, it can be accounted for in the timeseries through temporal interpolation as described.

#### Annotations

Words and faces were annotated in the movies so that they could be used to show data quality and timing accuracy while also illustrating a fairly straightforward method to label brain networks. To be used for this purpose, the overall quality of the annotations themselves needs to be demonstrated. For words, Table 3 provides a breakdown that reflects relative word on and offset accuracy for individual movies. Machine learning-based speech-to-text word transcriptions are assumed to have the highest temporal accuracy. An average of 75.75% of subtitle words had matching or similarity-matched word transcriptions. This was after hand transcribing over 2000 missing word times for ‘*Little Miss Sunshine*’ to bring accuracy up to 72.48% in order to correct for poor transcription accuracy (∼45%, possibly due to overlapping dialogue in the movie). Speech-to-text transcription left an average of 24.25% of the subtitle words to get estimated word lengths. Of these, an average of 20.30% were made up of the ‘continuous’ and ‘partial’ estimations, considered relatively accurate because they rely on accurately transcribed matched/similar words to make estimations. Only 3.95% of the subtitle words on average were fully estimated. These have the least accurate word timings because their length had to be estimated entirely from the subtitles page start and end times. Finally, to increase accuracy we truncated the 2.52% of words that were unreasonably long. In summary, it might be argued that about (75.75% Matched/Similar + 20.30% Continuous/Partial) = ∼95% of words have relatively accurate millisecond level onset times. Given that there are >10,000 words on average per movie, a ∼5% rate for less accurate word timing is likely acceptable.

**Table 3.**
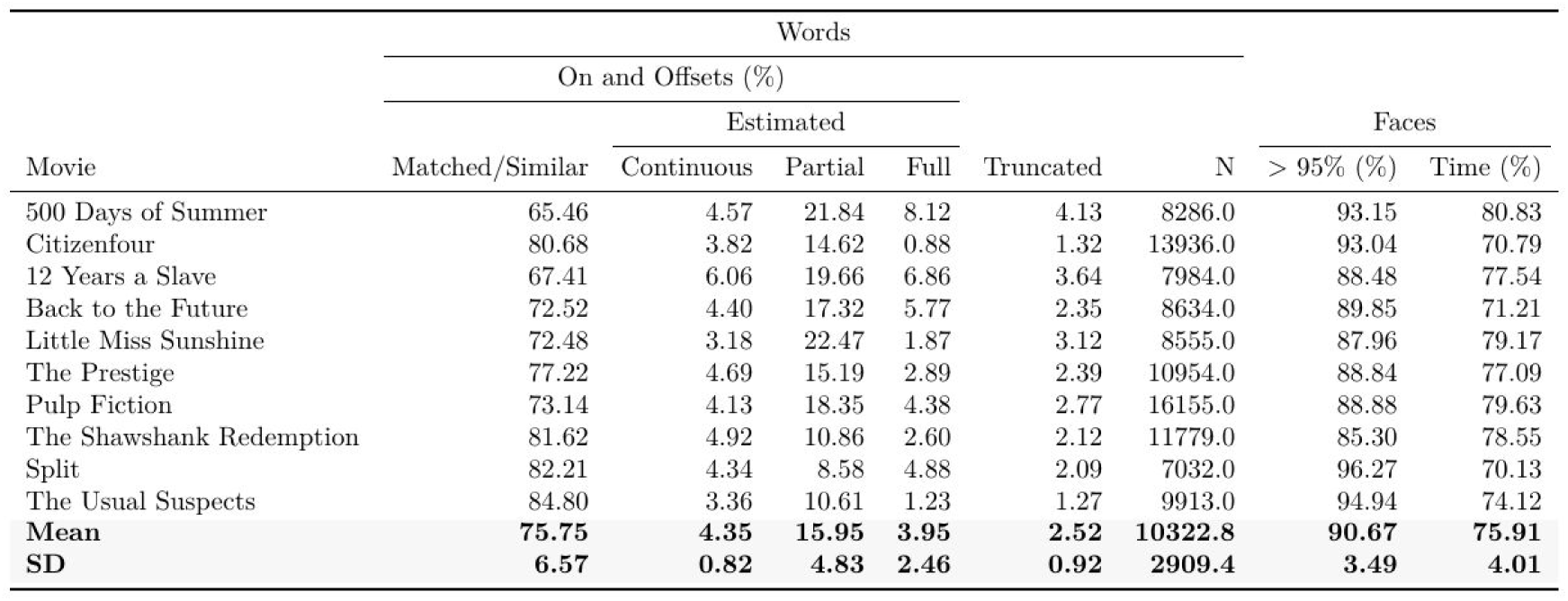
Movie word and face annotation information. The on and offsets of words were obtained from machine learning-based speech-to-text transcriptions. Dynamic time warping was used to align these to subtitles. If words in a subtitle page ‘Matched’ or were ‘Similar’ to words in the transcript, it received the transcript timing. Otherwise it was estimated. ‘Continuous’ estimations are single subtitle words inheriting the start and end time from the end of the prior and start of the next transcribed word. ‘Partial’ estimations are similar but involve two or more missing words between transcribed words. ‘Full’ estimations occured when no words were transcribed and words were estimated from the start and end time of the subtitle page. When word lengths were unreasonable, they were ‘Truncated’. This procedure resulted in an average number (‘N’) of >10,000 words per movie. The on and offsets of faces were also obtained from a machine learning-based approach. The final two columns are the average percentage of face labels with >95% confidence and the percent of time faces were on screen.

For face labels, histograms for all movies were used to examine the distribution of confidence levels. Across all movies, the average percentage of face labels with a confidence value greater than .95 was 90.67%, motivating us to use all the labelled faces in further analysis (Table 3). We also qualitatively compared results with the movies and they appeared to confirm that confidence levels were accurate.

### Anatomical MRI

Table 4 provides a list of anatomical and functional MRI irregularities. Anatomical image segmentation and cortical surface reconstruction with Freesurfer finished without error for all participants. Surfaces were individually inspected and no manual corrections were needed, suggesting anatomical images were of good quality.

**Table 4.** Data acquisition irregularities that might have impacted data quality. Most irregularities centred around participant drowsiness. We monitored participants through a camera and occasionally gave them warnings if they appeared drowsy to us. In a few cases we paused the scan to let participants compose themselves and to make sure they would remain alert throughout the rest of the scan.

| BIDS ID | Movie | Irregularity |  |
| --- | --- | --- | --- |
|  |  | Hardware / Scanning | Participant |
| 1 | 500 Days of Summer | Anatomical scan first | Paused to adjust volume |
| 14 | 500 Days of Summer | 30 channel headcoil |  |
| 16 | 500 Days of Summer | Given glasses at first break | Paused once b/c drowsy |
| 21 | Citizenfour |  | Paused once b/c drowsy |
| 24 | Citizenfour | Anatomical different day |  |
| 25 | Citizenfour |  | Appeared drowsy |
| 28 | Citizenfour |  | Paused b/c fell asleep once |
| 33 | Citizenfour |  | Appeared drowsy |
| 35 | Citizenfour |  | Appeared drowsy |
| 37 | Citizenfour |  | Paused once b/c drowsy |
| 38 | Citizenfour |  | Paused once b/c drowsy |
| 47 | Pulp Fiction | Paused to adjust earphones |  |
| 51 | The Shawshank Redemption | Wrong EPI sequence (2.8 s) |  |
| 55 | The Shawshank Redemption | Paused to clean glasses |  |
| 59 | The Prestige |  | Paused once b/c drowsy |
| 63 | Back to the Future |  | Appeared drowsy |
| 65 | Back to the Future |  | Paused to adjust volume |
| 73 | Split | Arduino issue / extra pauses |  |
| 80 | Little Miss Sunshine |  | Paused once b/c drowsy |

### Functional MRI

#### Temporal Signal-to-Noise Ratio

Mean tSNR for timeseries averaged over grey matter voxels was comparable to prior multiband four studies reviewed earlier. Furthermore, there were comparable increases in tSNR after preprocessing (Table 5). Cohen’s d at the individual voxel level shows regions of the brain for which tSNR increased after full preprocessing (Figure 2). This includes most medial and posterior aspects of the brain, with less tSNR increase in the frontal lobe.

**Figure 2.**
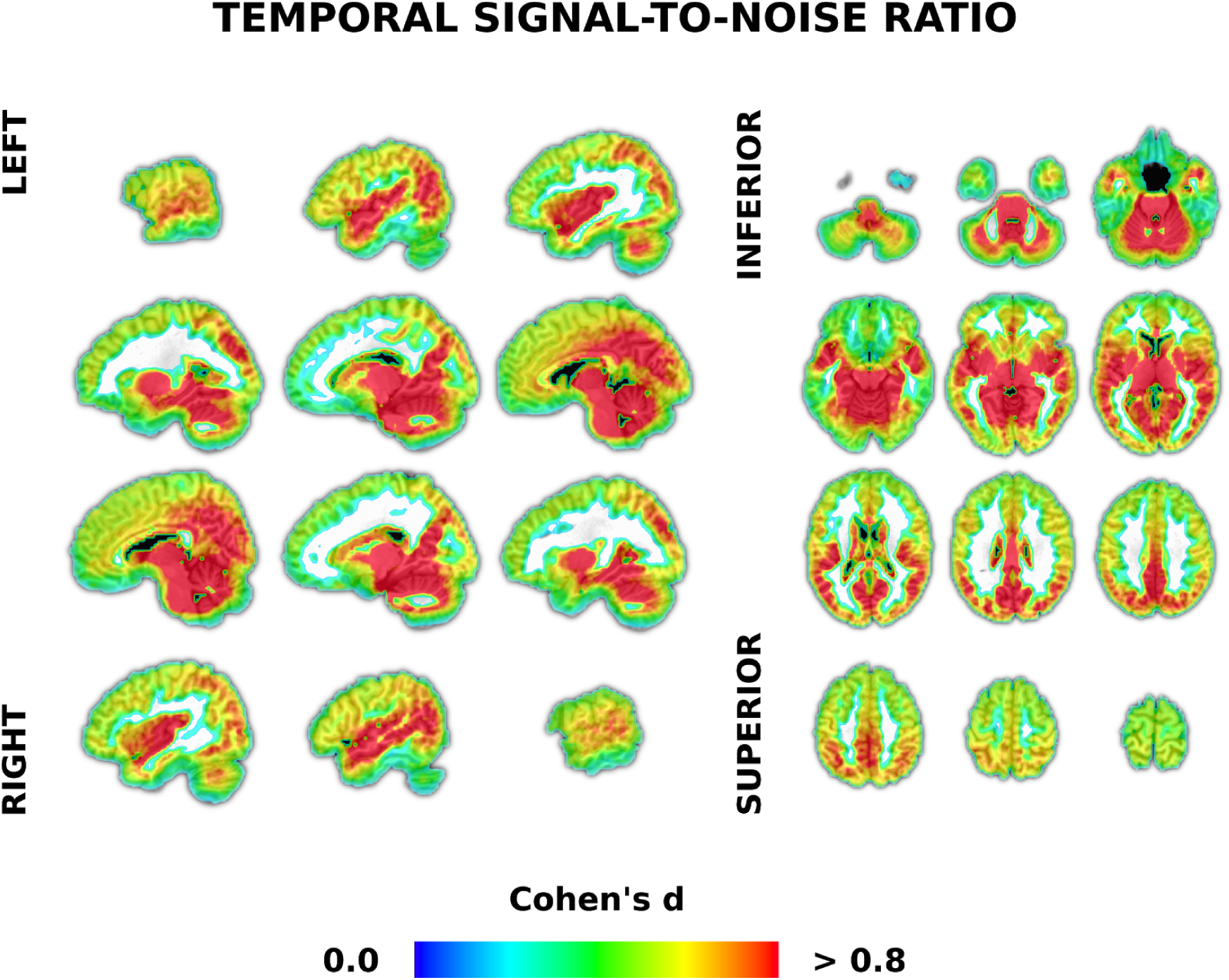
Voxel-wise temporal signal-to-noise ratio analysis demonstrating increases in data quality with preprocessing. Temporal SNR was calculated in each voxel using mostly unprocessed and fully preprocessed functional magnetic resonance imaging (fMRI) timeseries data from 86 participants. Full preprocessing included blurring and detrending using motion, white matter, cerebral spinal fluid and independent component analysis (ICA) based artifact regressors. Cohen’s d effect sizes were calculated in each voxel as the mean differences between fully preprocessed and minimally preprocessed fMRI timeseries tSNR, divided by the pooled standard deviation. See Table 5 for tSNR values averaged across grey matter voxels.

**Table 5.** Descriptive statistics of temporal signal-to-noise ratio and independent component analysis based measures of data quality across movies. The temporal signal-to-noise ratio (tSNR) was calculated in mostly grey matter for minimally preprocessed (‘Min Pre’), blurred (‘Blur Pre’) and fully detrended and preprocessed (‘Full Pre’) functional magnetic resonance imaging timeseries data (see also Figure 2). The final column is the percent of manually-labelled independent component analysis (ICA) artifacts out of 250 dimensions.

| Measure | tSNR |  |  |  |  |  | ICA artifacts (%) |
| --- | --- | --- | --- | --- | --- | --- | --- |
|  | Mean |  |  | Maximum |  |  |  |
|  | Min Pre | Blur Pre | Full Pre | Min Pre | Blur Pre | Full Pre |  |
| Min | 11.85 | 12.20 | 13.37 | 91.33 | 110.33 | 185.85 | 56.40 |
| Mean | 39.43 | 44.55 | 63.82 | 161.19 | 201.20 | 319.18 | 71.71 |
| SD | 10.17 | 12.73 | 20.79 | 29.15 | 40.16 | 50.58 | 7.47 |
| Max | 60.10 | 68.99 | 98.03 | 218.09 | 310.91 | 431.79 | 87.20 |

#### Overall Data Quality

We assessed overall fMRI timeseries data quality using 10 measures and a composite of these. Table 6 shows the means per run across participants for eight of these measures. With the exception of run three, the number of outlying timepoints was under 1% per run on average. Maximum motion, as measured by six motion regressors, was low, under a degree and millimeter on average. The greatest maximal displacement was pitch (0.96°) and movement in the inferior/superior direction (0.92 mm). This is perhaps what might be expected for supine participants whose heads are firmly held in the left/right directions. Maximum delta was similarly under one millimeter. These parameters did not increase more in later compared to earlier runs. If anything, maximal movement decreased over the scanning session. The two other measures of overall data quality are given in Table 5, showing that tSNR (discussed in the prior section) and the number of ICA artifacts were reasonably high and low, respectively. On a subset of 50 datasets, there was a 96.22% agreement (SD = 2.20) between authors with regard to ICA artifact classification. Percentages of ICA artifacts are similar to those found in prior studies reviewed earlier. Finally, we created a composite measure from these 10 metrics to detect outliers (reverse coding tSNR). These measures had a high internal consistency with Cronbach’s alpha = 0.94. Using this measure, only one participant was considered an outlier. This is the participant mentioned in the Methods/Participants that was excluded from the database. Taken together, these measures indicate that NNDb timeseries data are of high overall quality.

**Table 6.**
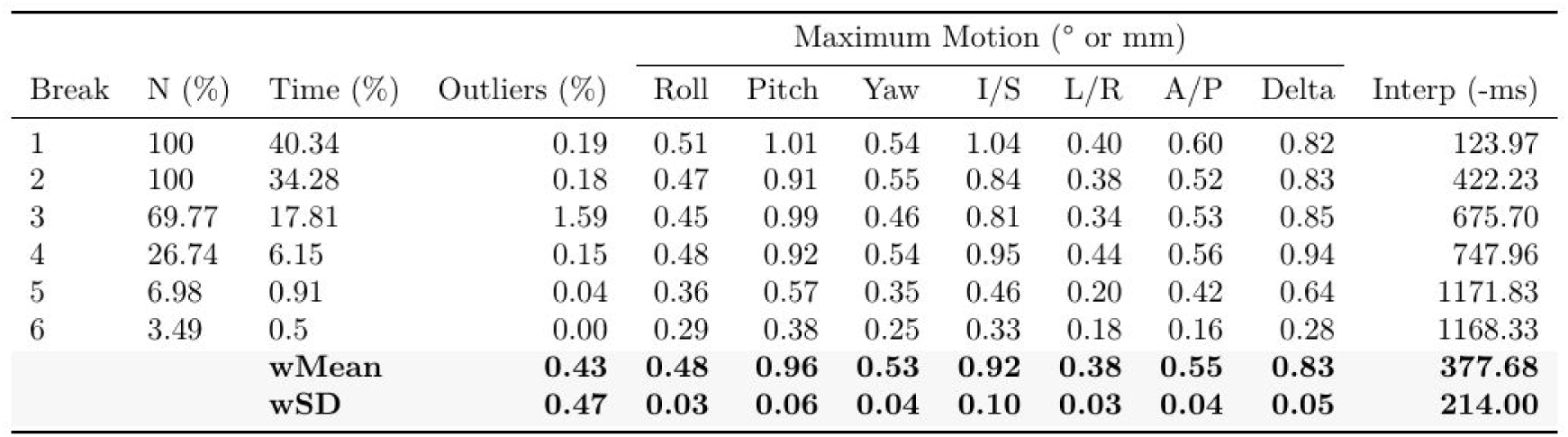
Descriptive statistics for outlying timepoints, motion and timing measures of data quality averaged over movie runs. ‘N’ the percentage of 86 participants having up to six breaks during any given movie. ‘Time’ is the average percentage of the whole movie for the run preceding each break. ‘Outliers’ is the mean percentage of timepoints with greater than 10% outliers in each run. Motion includes the mean maximum deflection in the inferior/superior (‘I/S’), left/right (‘L/R’) and anterior/posterior (‘A/P’) directions and the mean maximum change between any two timepoints (‘Delta’) in millimeters (mm). ‘Interp’ is the amount timeseries were interpolated back in time in milliseconds (-ms) in each run on average to account for known delays. The bottom two rows are the weighted means (wMean) and standard deviations (wSD) of rows weighted by the Time column.

#### Intersubject Correlation

ISC was done to show functional fMRI data quality and timing accuracy by demonstrating synchronization between participants and movies. There was significantly higher ISC at a Bonferroni-corrected threshold in large portions of auditory and visual cortices (precisely following sulci and gyri) when participants watched the same movies compared to different movies (Figure 3, top). Similar to prior work, the maximum correlation was *r* = 0.28. To examine reliability, we split the movies into two groups of participants that watched different sets of five movies. The results were largely spatially indistinguishable from each other (*r* = 0.96) or from results with all movies (with *r*s of 0.991 and 0.987). We also calculated spatial overlap using the Dice coefficient (or the Sorensen-Dice index) on data thresholded at a *t*-value of 10 (Figure 3, bottom). This was an arbitrary value chosen because even extremely high p-values resulted in whole-brain ISC. The resulting Dice coefficient was 0.82. These results demonstrate high data quality through robust activity patterns, spatial precision and timing accuracy through participant synchrony with movies.

**Figure 3.**
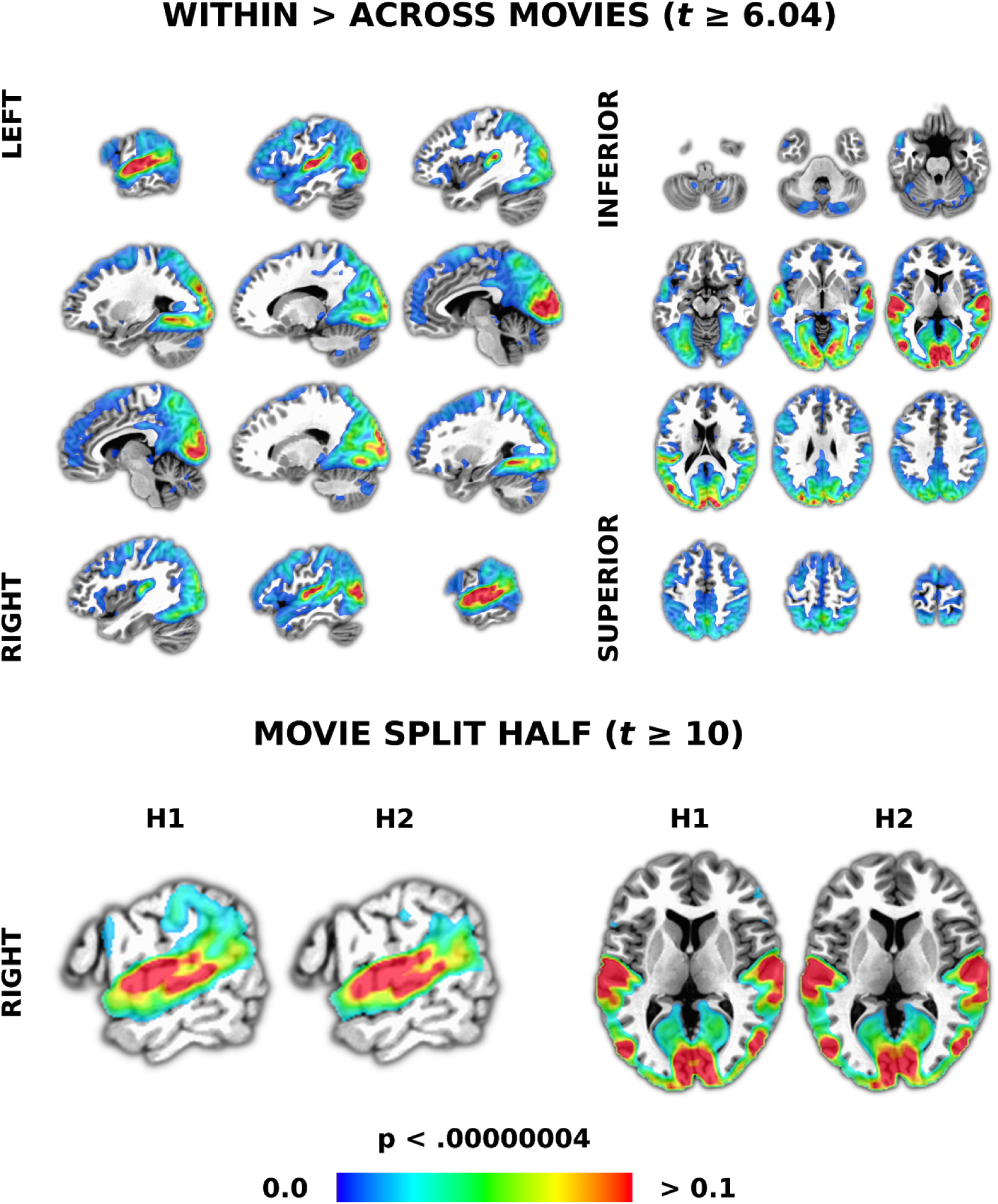
Results of intersubject correlation (ISC) demonstrating data quality and timing synchrony between participants and movies. ISC is a data-driven approach that starts with calculating the pairwise correlations between all voxels in each pair of participants. We used a linear mixed effects with crossed random effects (LME-CRE) model to contrast participants watching the same versus different movies (top). Equally-spaced slices were chosen to be representative of results across the whole brain. To demonstrate reliability, we split the data in half, with each having five different movies. The same LME-CRE model was run on each half and the results are presented at an arbitrary threshold to more easily view similarities and differences (bottom row). Slices were chosen to make differences more salient. The colour bar represents correlation values (r) in all panels. All results are presented at a p-value corrected for multiple comparisons using a Bonforoni correction and an arbitrary minimum cluster size threshold of 20 voxels.

#### Network Labelling

ICA and regression with a canonical response function were used to demonstrate data quality, timing accuracy and an approach to network labelling. There were M = 11.43 (SD = 4.31) word > no word, M = 13.52 (SD = 7.21) no word > word, M = 8.71 SD = 8.09, face > no face and M = 8.44 (SD = 7.84) no face > face networks per participant, each significant at a stringent Bonferroni-corrected threshold. For words (compared to no words), these networks variously consisted of activity in the superior temporal plane, posterior inferior frontal gyrus and motor regions as might be expected during language processing^89^. For faces (compared to no faces), activity was in the posterior superior temporal sulcus and fusiform gyrus among other regions that might be expected during face processing. An example from a single participant is shown in Figure 4 (top), using hierarchical clustering (with Ward’s method) to order all significant word > no word and face > no face networks in terms of the Euclidean distance between IC timecources to show network similarity. By this approach language and face networks mostly cluster separately.

**Figure 4.**
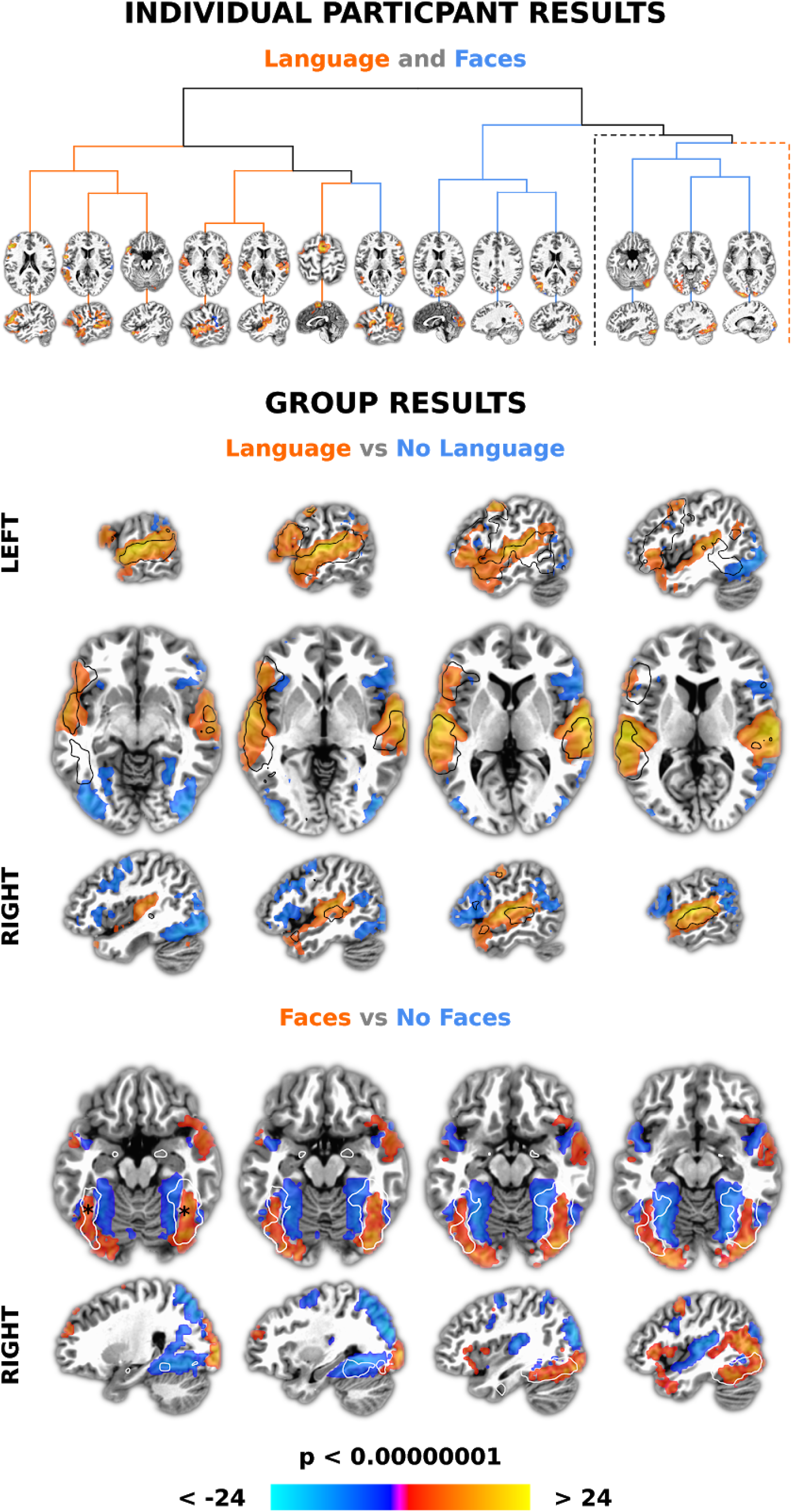
Results of combined independent component analysis (ICA) and model-based analysis demonstrating data quality, timing accuracy and an approach to network labelling. First, networks were found at the individual participant level using ICA, a multivariate data-driven approach. Word and face annotations from movies were then convolved with a standard hemodynamic response function and used in general linear models to find associated IC timecourses. The dendrogram (top) shows 13 of 20 significant networks from an example participant that were more associated with words > no words (‘Language’; red lines) and faces no faces (‘Faces’; blue lines), clustered to show IC timecourse similarity. Slices are centred around the centre of mass of the largest cluster in each network. Two branches (dotted lines) were excluded for visibility. These had an additional five language and two face networks. For group analysis, spatial components corresponding to significant IC timecourses for each participant were summed and entered into *t*-tests. The middle panel shows that word > no word networks (‘Language’; reds) overlap a ‘language’ meta-analysis (black outline) more than no word > word networks (‘No Language’; blues). Slices are centred around the centres of mass of the two largest clusters, in the left and right superior temporal plane. The bottom panel shows that face > no face networks (‘Faces’; reds) produced greater activity than no face face networks (‘No Faces’; blues) in the same areas as a ‘fusiform face’ area (FFA) meta-analysis (white outline). Slices are centred near the average x/y/z coordinates of the putative left and right FFA (indicated with black asterisks). The colour bar represents z-scores in all panels. All individual and group level results were Bonforoni corrected for multiple comparisons and presented with an arbitrary minimum cluster size of 20 voxels.

Group t-tests across participants showed significant patterns of activity consistent with the sum of networks from individual participants, again using a Bonferroni corrected threshold. In particular, word and face networks resembled meta-analyses of language (Figure 4, middle, black outline) and face processing (Figure 4, bottom, white outline). The face networks included the putative fusiform face area(s) (Figure 4, bottom, black asterisks), with immediately adjacent regions more involved in processing times in the movie when faces are not visible. Overall, both individual and group results demonstrate high data quality by being robust and showing anatomical precision. Furthermore, such strong relationships between stimulus annotations and idealised timeseries again indicate that timing accuracy is high.

## Usage Notes

We think that the NNDb has the potential to help revolutionise our understanding of the complex network organization of the human brain as it functions in the real-world. However, there are several usage bottlenecks, including annotations and analyses that we now discuss to help others use the NNDb to make new discoveries. We then conclude by briefly discussing how collective community based annotations and usage will contribute to the future of the NNDb.

### Annotation Bottleneck

Annotations are necessary for hypothesis testing using the NNDb. This involves not only coding a stimulus feature of interest but also a suitable range of controls at a finer level of detail than used to label ‘language’ and ‘face’ networks herein. For example, if one were interested in the neurobiological mechanisms of how observed face movements are used by the brain during speech perception^90,91^, one might want to annotate a large range of features. These might include speech with more or less environmental noise when the face is visible (as audiovisual speech improves speech in noise). There might need to be annotated auditory-only controls matched for auditory/semantic features and visible scene complexity. There may need to be face-only controls or audiovisual controls with faces in profile, etc. If done manually, movie annotations at this level of detail might be very time consuming. Though this might prove necessary for testing some specific hypotheses, we suggest automated approaches and a brain-driven approach that might be used to speed up the annotation process.

Automated approaches to annotation can make use of a large number of existing text-based descriptions of movies to provide time-locked features. These include, 1) Detailed descriptions in scripts that can be aligned to movie times from subtitles^92^; 2) Detailed verbal descriptions from descriptive video services that make movies accessible to millions of visually impaired and blind individuals^93^; 3) Video clips from movies available on social websites like YouTube that can be matched to movie times by visual scene matching to include user comments as features^94^. For example, one two-minute clip from Gravity on YouTube currently contains over 3,800 comments that can be text-mined for features; and 4) There are many emerging automated machine learning approaches for labelling features, e.g., the YouTube8M which has a vocabulary consisting of 4716 features (e.g., ‘cat’, ‘book’, ‘egg’, etc)^95^ or human action video datasets^96^.

Brain-driven annotations potentially decrease the need to annotate everything in movies. That is, the brain data itself can be used to identify movie timecodes for acquiring more detailed annotations. This allows users of the NNDb to focus on times when networks of interest are processing information, reducing the amount of movie that needs to be annotated. For example, ICA can be used to derive networks and associated independent component timecourses (as shown in Figure 4, top). Users of the NNDb can annotate only what happens when the response is rising (or at peaks) in these timecourses in components that represent networks of interest (thus, being able to determine what the 11 individual participant networks grossly labelled as ‘Language’ are doing in Figure 4). This can be done manually, with the aforementioned automated approaches or in a crowdsourced manner. For example, one could submit the videos from the at rise times in IC timecourses in Figure 4 (top) containing the amygdala and have thousands of people quickly label observed emotional characteristics.

### Analysis Bottleneck

Another potential bottleneck is analysis. There are arguably no standardised approaches for analysing complex and high dimensional fMRI data from long natural stimuli like movies (though a few approaches are becoming increasingly common^58,97,98^). The computer science community has learned that an extremely effective way to foster research on a topic is by running machine learning competitions on fixed datasets. These competitions allow unambiguous comparison of solutions to a problem and allow small improvements to be clearly noted and published. For example, the annual ‘ImageNet Large Scale Visual Recognition Challenge’ (ILSVRC) resulted in algorithms that outperform humans far more quickly than expected^99^. Machine learning approaches are becoming an increasingly common way to analyze fMRI data, with a growing number of examples applied to natural movie stimuli^98,100^. We suggest that, to generate innovation in analysis, that competitions similar to the ILSVRC could be run using the NNDb to crowdsource the development of new machine learning (and other) approaches for fMRI data from movies.

### Future of the NNDb

We will make more fMRI data available as it is acquired at the same (1.5T) and higher field strengths (3T) in typically developing and clinical groups. We hope that over time, we will be able to amass the collective effort of our own and other communities to collate annotated stimulus features to be able to ask more and more specific questions of the data. We will make these annotations and improvements to older annotations available in regular updates. We will make code for analysis available, e.g., we are developing a graph theoretic dynamic core-periphery algorithm capable of fast voxel-based analysis. We will try and host machine learning-based analysis competitions to drive innovation and make all code available.

## Code Availability

Scripts used in this manuscript are available at: https://github.com/lab-lab/nndb_project. Additional information can be found at http://naturalistic-neuroimaging-database.org/.

## Acknowledgements

This work was partially supported by EPSRC EP/M026965/1 (JIS) and doctoral training awards from the EPSRC (FG) and BBSRC (SA). We would like to thank Fred Dick, Tessa Dekker, Joe Devlin and other staff at the Birkbeck-UCL Centre for Neuroimaging (BUCNI) for an incredible level of support. Thanks to Abbas Heydari for creating the first version of the Arduino/movie pausing solution. Thanks to LAB Lab stallions who helped with the NNDb: Dyutiman Mukhopadhyay, Alberto Arletti, Letitia Ho, Ellie Pinkerton, Yixiao Wang and Pippa Watson. JIS would like to thank his Banana for support. Thanks to Daniel Lametti, Peter ‘PBR’ Kirk and Dyutiman Mukhopadhyay for comments on the first draft. We thank Gang Chen at the NIH for statistical advice and help implementing the LME-CRE analyses.

## Author Contributions

JIS conceived the NNDb and wrote the manuscript with help from all authors. All except JIS and SM collected data, led by SA as operator. JIS and SA did the preprocessing and technical validation of the f/MRI data. FG wrote the movie pausing script and created face annotations. JIS and FG developed the temporal interpolation procedure. JH transcribed the questionnaires and created word annotation scripts. SA and JIS did the tSNR analysis. SM did the ISC analysis. JIS did the network labelling analysis.

## Supplementary Material

We supplement the main text with further information on how we derived word annotations from the movies, how we paused movies and subsequently corrected for timing delays caused by pausing.

### Word Annotations

Here we provide further technical information on how word annotations were generated. First, to find subtitle and transcription ‘matches’ and similarity matches:

- Dynamic time warping (DTW) was used to find the alignment between each subtitle page and transcribed words, starting 0.5 seconds before and ending 0.5 seconds after the page to account for possible subtitle inaccuracies.
- All punctuation was removed from the subtitles.
- To better match words with partially accurate transcriptions, the transcripts and subtitles were stemmed (e.g., ‘kittens’ becomes ‘kitten’). Once the word match checking was complete, the words were restored to their original unstemmed form.
- If one subtitle word aligned with one or more transcription words and there was a match between those, we use the transcription timing of the word that matches the subtitle word.
- If the words were not the same, we used the timing of the transcription word with maximum Jaro similarity if Jaro similarity was > .60 with that subtitle word.
- If multiple subtitle words aligned with one transcription word (e.g., ‘is’,’’a’, ‘story’ in the subtitles and ‘story’ in the transcription), we gave the timing of the transcribed word to the matched subtitle or most similar word if the Jaro similarity was > .60.

Next, for subtitle words that have no match and max similarity < .60, or in case of multiple subtitle words match one transcript word (e.g., ‘this’, ‘this’, ‘is’ in the subtitle and ‘this’ in the transcript), we estimate the timing of these words using the following methods:

- A container was used to include words whose timings werte not obtained from a direct or similarity match.
- Once matched/similar words are found, we estimate the start times of the words in the container based on a previous known timepoint, usually the end time of the previously matched/similar word, or the start time of the subtitle screen. The end time is based on the start time of the next matched/similar word.
- At the end of a subtitle page, if the start time of the next line is one second longer than the end time of the current line, we estimated timing with the end time being the end time of the subtitle screen.
- Depending on the number of words in the container when making estimations, they are labelled as ‘continuous’ or ‘partial’:
  - If there is only one word, the word is labelled as ‘continuous’, with the assumption that the timing for the word should be reasonably accurate if it is between two words in continuous speech.
  - If there are multiple words, the words are labelled as ‘partial’, and the start time and end time of each word is estimated based on the number of letters in each word (as described in the manuscript).
- If the subtitle is not accurate, the start time will be later than the end time for the words not estimated (i.e., the start time of the subtitle screen is later than start time of a matched word). Here, we estimated words based just on end times, assuming each word has a duration equal to the number of letters times 0.03 seconds.

If no words are transcribed in that window, all the words in the subtitle page are estimated, based on the start and end time of the subtitle page and the number of letters in each word (as described in the manuscript). These words are labelled as ‘full’ estimations. Finally, at the end of these steps, the script does some post-processing:

- We reordered words based on onset times, removing words with the same timings.
- If words overlapped, we shifted the start time of the word to the end time of previous words.
- For numbers (e.g. 32) not correctly identified in the transcription, we changed to the spelled form (‘thirty two’) and re-ran the script.
- We truncated unusually long words. For example, two four letter words in a 10 second window would each be estimated as 5 seconds long. As this is unreasonable, we truncated estimated words < 10 letters and more than 2.5 standard deviations from the mean word length in conversational speech to the mean (based on^65^). Specifically:
  - Words more than 1000 ms and < 10 letters in length were truncated to 600 ms.
  - As it is common for words more than 10 letters to be longer than 1 second when spoken, estimated word lengths for words with >10 letters and < 2 seconds were kept
  - Estimations > 2 seconds were truncated to 1000 ms.

### Movie Pausing

Here we provide additional information about how the movie pausing script/hardware solution kept track of timing so it could subsequently be used to rewind movies and to account for timeseries delays in preprocessing.

Whenever the scanner was stopped and a movie was paused, the whole brain volume or TR being collected was dropped. In the first version of our script, the movie simply restarted where it had been stopped when scanning was resumed. In the second version, we edited the script to account for the dropped volume by rewinding the movie. To calculate the amount of movie time lost since the beginning of the dropped volume, the script uses three output files it generates when running: A *MPlayer output* file, *current time* file and *final output* file (all in ‘.txt’ format).

The role of the *MPlayer output* file was to enable the script to read the current time position in the movie. Every time the BASH script checked for a new TTL pulse (i.e. every 50ms), it would also send a command to MPlayer to get the time position in the movie (using the *pausing_keep_force* and *get_time_pos* commands for MPlayer in slave mode). As MPlayer received commands through a temporary */tmp/doo* file, the script had to pipe the stdout output to the *MPlayer output* file for it to then be able to read the value itself. MPlayer only gives the time position up to one decimal. A line inside *MPlayer output* would look like:

ANS_TIME_POSITION=1.6

The script would then read the last line of the *MPlayer output* file and write a new line in the *current time* file. Every line consisted of the newly acquired time position in the movie and a timestamp formed by the Linux epoch time (the number of seconds from 00:00:00 UTC on 1 January 1970) and the milliseconds elapsed since the end of the previous second. A line inside the *current time* file would look like:

1572708345 209 ANS_TIME_POSITION=1.6

If paused, the movie is then rewound by that amount by passing a command to Mplayer through ‘slave’ mode. When the scanner is restarted, the movie begins within 100 ms of the first TTL pulse (again, because it had to monitor at least two pulses). Because of a coding error, version two of the script occasionally fast forwarded when it should rewind. After fixing this error, the movies rewound correctly whenever the scanner was stopped for the remaining participants. In all versions, when a TTL pulse is received, signalling that the scan has restarted, eight seconds of discarded acquisitions are acquired before the movie is unpaused.

Whenever the movie was paused or started, the script would write to the *final output*, which would typically contain the following lines:

1567528264 953 start

1567531380 437 pause 1

1567531465 886 rewind -.592 start

1567534037 162 pause 2

1567534091 303 rewind -.384 start

1567535208 234 ended

The above example is taken from the last version of the script, which included the rewind values. The first version did not include these values. To calculate the rewind times, the script would read the last *start* and *pause* lines of the *final output* file:

1567528264 953 start

1567531380 437 pause 1

Because our TR=1s, we started counting the number of total TRs registered from the timestamp of the start value in *final output*. For example, above we would consider 3116 TRs elapsed from 1567528264 953 until 1567531380 953 (1567531380 - 1567528264). However, as the script stopped the movie at 1567531380 437 only 3115 TRs were registered, meaning that the registered data only went up to 1567531379 953. So, the number of milliseconds of the movie playing without any brain data being acquired would be the difference between 1567531379 953 and 1567531380 437, which would be 437 + 1000 - 953 + 108 = 592. The 108 value was added to account for the fact that it would actually take 108 ms from the moment the script registers the start of a new TR and when the play command is given to MPlayer (the script would pause for 100ms, while the other 8ms delay was observed during piloting and assumed to be due to other running processes).

The reason behind the coding error in the second version of the script was a minus sign needed when the milliseconds in the pause time were greater than the milliseconds in the start time. The following example is from a correct working version of the script:

1561977334 281 start

1561980159 470 pause 1

1561980228 411 rewind -.297 start

There would be 2825 TRs registered between 1561977334 281 and 1561980159 281, leaving 470 - 281 = 189 milliseconds lost. The rewind time would be 189 + 108 = 297ms, with a command being sent with a minus sign in front (a lack of a minus sign would fast forward by that amount of ms). To distinguish between the two cases an if statement was used. However, in the second version of the script the minus sign was accidentally omitted in one of the branches of the script, resulting in the error described. The *current time* and the *final output* files were later used as inputs into a Python script that calculated the number of milliseconds lost each time the movie was paused.

### Timing Correction

Here we describe how we account for the delays generated by the timing scripts. Specifically, when the scanner is stopped, the TR being collected is dropped. Thus, some portion of the movie is played but there is no corresponding TR in the timeseries. In v1 of the mplayer script, that time is lost (because the movie was not rewound). To account for this, the TR is added back to the run. This is done so by retrieving the last timepoint of the run in which the movie was stopped and the first timepoint of the run after the movie was stopped and averaging these. This makes the transition between TRs less abrupt.

Next we shift the timeseries through interpolation (using ‘*3dTshift*’). In version one of the MPlayer script, the amount of time dropped in run one will determine how far back to interpolate run two and so on. For example, if the movie stopped at 1000.850 and the last full TR was lost, it means that 850 ms of the movie was watched but is missing from the timeseries. To account for the missing information, we add a TR to the timeseries being collected before the scanner was stopped (done above) and interpolate the next run backwards in time the amount not covered by this TR. Thus, for the 850 ms of movie not watched, this means there is 150 ms too much time added to the movie by adding a TR (because our TR = 1 second). So we shift the next run back this amount so that the timeseries is theoretically continuous again (though this is never really possible). If there is another run (i.e., three or more), the same logic applies except that the extra 150 ms needs to be accounted for. So, if the next run stopped at 2000.900, we shift run three back (1000-900)+150 ms = 250 ms. These calculations are complicated by the fact that each scanner stop is always a 100 ms delay and a known standard deviation because of the way the MPlayer script works (see above). For this reason, every run is time shifted backward this extra amount. So in the example, if this delay was 100 ms, run three in the prior example would be shifted back 350 ms.

Version two of the script is simpler. That is, we rewound the movie the exact amount that we lost when the scanner stopped. Thus, an additional TR is not added and the only time shifting corresponds to the time lost whenever the scanner was stopped from monitoring for the TTL pulse. Unfortunately, an error in the script initially caused some of the runs to occasionally be fast forwarded. In these cases, the timing correction was carried out as in the prior paragraph. In all other cases, the rewind feature assured that cumulative delay is the only time shifting. For example, if there are three runs and 100 ms was lost each run, the final run would be time shifted back 300 ms.

The timing for one dataset was further corrected due to technical issues with the Arduino device wires on the day of the scan. The Arduino mistakenly stopped transmitting the TTL pulse, likely because of a loose wire, registered by the BASH script as pauses when the scan was still ongoing. Thus, instead of the two actual pauses, eight were recorded, meaning six of the alleged pauses did not occur. The false pauses added eight seconds to the timing output file as the scan was still ongoing, increasing the apparent total length of the movie by 48 seconds, and therefore increasing scan time as a consequence. In order to correct for this error, eight TRs were removed from the timeseries whenever a false pause was detected, for a total of 48 TRs removed.

